# mRNA structure determines specificity of a polyQ-driven phase separation

**DOI:** 10.1101/233817

**Authors:** Erin M. Langdon, Yupeng Qiu, Amirhossein Ghanbari Niaki, Grace A. McLaughlin, Chase Weidmann, Therese M. Gerbich, Jean A. Smith, John M. Crutchley, Christina M. Termini, Kevin M. Weeks, Sua Myong, Amy S. Gladfelter

## Abstract

RNA promotes liquid-liquid phase separation (LLPS) to build membrane-less compartments in cells. How distinct molecular compositions are established and maintained in these liquid compartments is unknown. Here we report that secondary structure allows mRNAs to self-associate and determines if an mRNA is recruited to or excluded from liquid compartments. The polyQ-protein Whi3 induces conformational changes in RNA structure and generates distinct molecular fluctuations depending on the RNA sequence. These data support a model in which structure-based, RNA-RNA interactions promote assembly of distinct droplets and protein-driven, conformational dynamics of the RNA maintain this identity. Thus, the shape of RNA can promote the formation and coexistence of the diverse array of RNA-rich liquid compartments found in a single cell.

**One Sentence Summary:** Identity in cellular, phase-separated compartments arises from RNA-RNA complexes encoded by mRNA secondary structures.

## Main Text

Formation of non-membrane bound organelles through the condensation of macromolecules is a recently appreciated mechanism of intracellular organization. These liquid-like condensates form through liquid-liquid phase separation (LLPS) and are found in the cytoplasm and nucleus (*1*, *2*). A fundamental unsolved problem is how liquid droplets recruit distinct constituents and retain independent identities. RNA can drive LLPS and modulates the material properties of droplets (*3*–*6*), but it is unknown if RNA controls the identity and maintenance of coexisting liquid compartments. Here we show mRNA secondary structure is required for droplet identity through directing interactions between mRNAs and RNA-binding proteins.

Whi3, a polyQ-containing, RNA-binding protein first identified in *Saccharomyces cerevisiae* (*7*), functions in morphogenesis, memory of mating, and stress responses, where it forms aggregates and associates with RNA-processing bodies (*8*–*11*). The homolog in the filamentous fungus *Ashbya gossypii* has one RNA recognition motif (RRM) and an expanded polyQ tract (Fig. S1A), and both regions promote self-assembly. In vitro, Whi3 polyQ-dependent LLPS is driven by specific RNAs encoding regulators of either the cell cycle (e.g. the cyclin *CLN3*) or actin (e.g. the formin *BNI1* and *SPA2*) (*3*). Distinct types of Whi3 droplets form in *Ashbya* cells: perinuclear *CLN3* droplets and *BNI1* droplets at sites of polarized growth at cell tips ((*12*, *13*), Fig. 1A and movie S1). These two types of droplets have different Whi3 levels and Whi3 incorporation rates (Fig. 1, B and C), suggesting they are structurally distinct.

**Figure 1.**
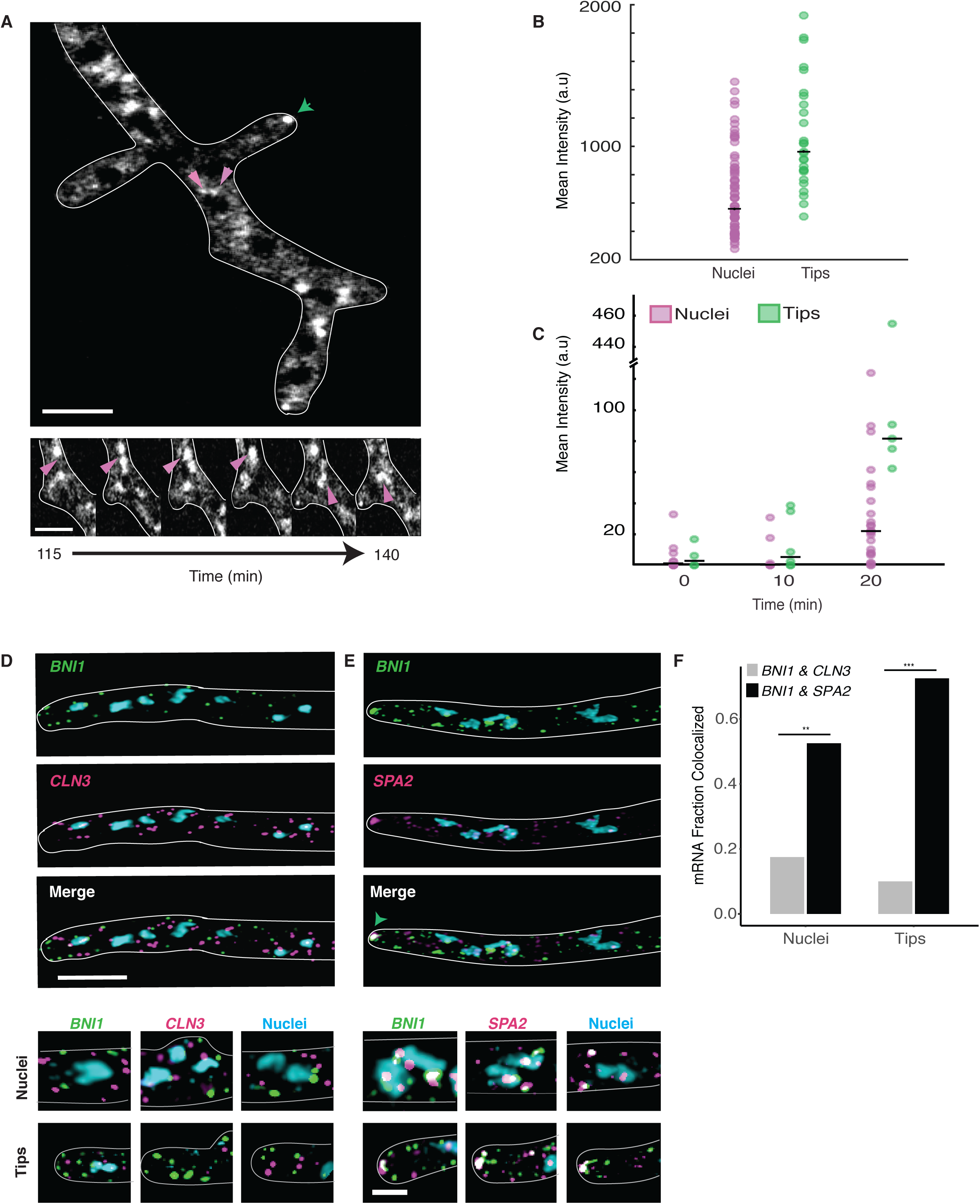
Cyclin and Polarity complexes are spatially and physically distinct within the cell. **A.** Top, Whi3 forms liquid droplets in *Ashbya gossypii*. Below, Whi3 droplets accumulate and fuse around nuclei. Green arrows denote polarity droplets. Pink arrows denote perinuclear droplets. Scale bars 5 μm. **B.** Mean intensity of Whi3-tomato is higher in tip droplets (green) than perinuclear droplets (pink). **C.** Rate of Whi3 incorporation is higher in tip compared to perinuclear droplets. **D.** smFISH images show *BNI1* (green) and *CLN3* (pink) mRNAs are spatially distinct. Nuclei are in blue. Scale bar 5 μm. **E.** smFISH images show *BNI1* (green) mRNAs co-localize with polarity mRNA *SPA2* (pink). Nuclei are in blue. The green arrow marks where the RNAs overlap at the tip. Inset scale bar 2 μm. **F.** *BNI1* and *SPA2* are significantly more co-localized than *BNI1* and *CLN3*. p<0.001 for tips and p<0.01 for nuclei (Fisher’s Exact test). n= 40 nuclei and tips for ≥30 cells.

The distinct droplet properties may depend on extrinsic features of the local cytosolic microenvironment or arise due to different droplet constituents. *CLN3* and *BNI1* mRNAs minimally co-localize in the cytoplasm by single molecule (sm) RNA F.I.S.H., although they were occasionally co-expressed by the same nucleus (Fig. 1, D and F). The lack of co-localization suggests there are intrinsic, compositional differences between droplets. In contrast, mRNA of the polarity regulator *SPA2*, frequently co-localized with *BNI1* mRNAs, especially at growth sites (Fig. 1, E and F). Thus, mRNAs encoding functionally related proteins co-localize, while functionally unrelated mRNAs do not. How can distinct Whi3-binding mRNAs segregate to different droplets in a common cytoplasm?

To address this question, we employed a reconstitution system to test if mRNA sequence was sufficient to generate droplet individuality (Fig. 2A). In vitro, as in cells, droplets composed of *BNI1* mRNA displayed higher Whi3 to RNA molar ratios than droplets made with *CLN3* mRNA (Fig. S1B). Remarkably, when *CLN3* mRNA was added to Whi3 droplets made with *BNI1* mRNA, *CLN3* preferentially assembled into new droplets, rather than incorporating into *BNI1* droplets (Fig. 2, B and C, S1C). In contrast, *BNI1* mRNA readily incorporated into preformed droplets (Fig. 2, B and C). Notably, *SPA2* mRNA incorporated into *BNI1* droplets (Fig. 2, B and C), and *CLN3* did not incorporate into *SPA2* droplets (Fig. S1D). Thus, as in cells, cyclin and polarity mRNAs assemble into distinct and immiscible droplets in vitro, indicating droplet identity is encoded by the mRNA.

**Figure 2.**
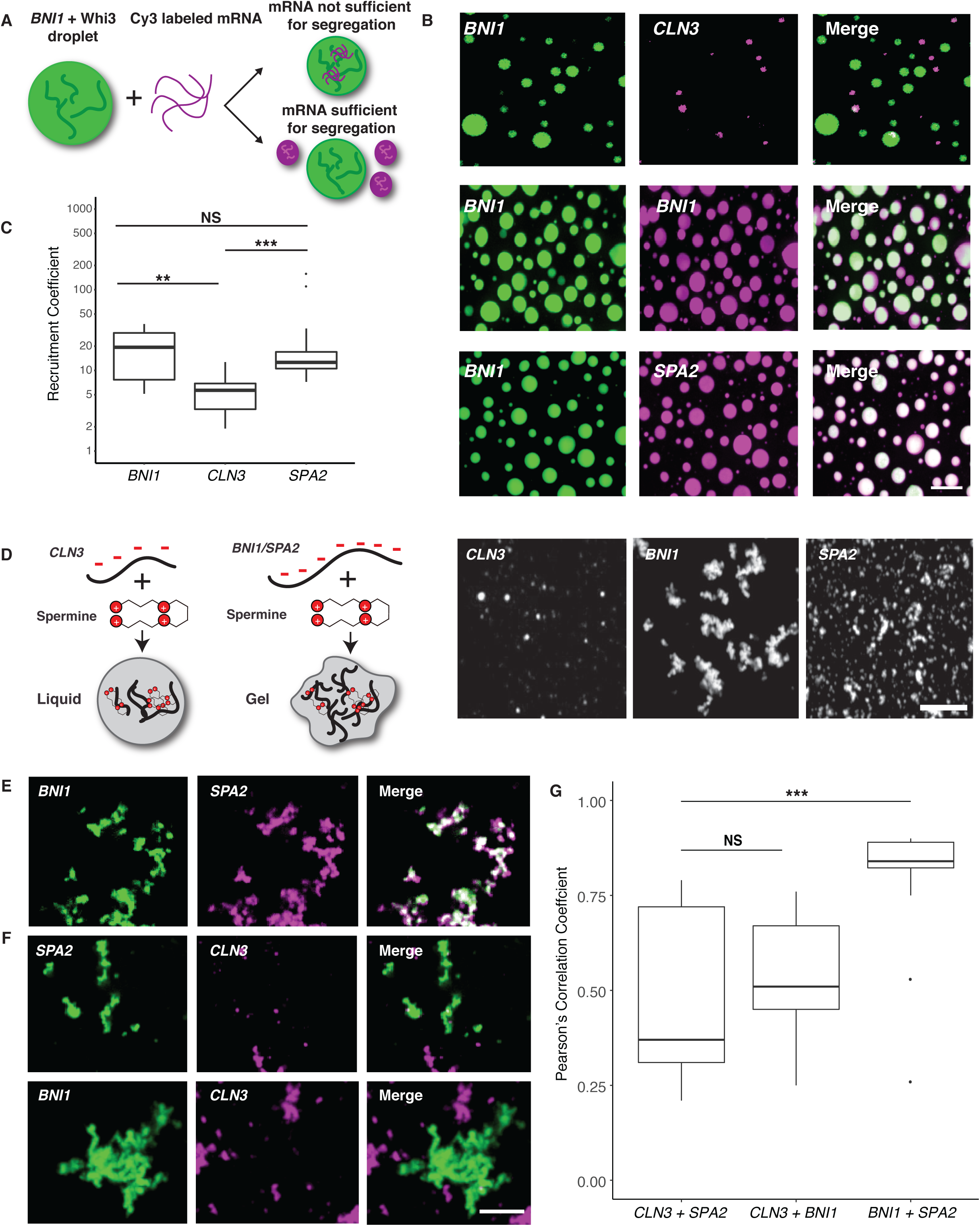
Polarity and cyclin complexes segregate *in vitro*. **A.** Experimental schematic of in vitro droplet recruitment assay. **B**. *CLN3* mRNA (pink) is not efficiently recruited but *BNI1* or *SPA2* mRNA (pink) are recruited into preformed Whi3-*BNI1* droplets (green) based on fluorescence microscopy. Scale bar 10 μm. **C.** Recruitment coefficients of mRNA from **B**. Boxes indicate interquartile range, line is median and whiskers contain points within three times the interquartile range, and outliers are indicated with (*) marks. NS, not significant, p > 0.05; **, p < 0.01; ***, p < 0.001 (t test). n ≥500 droplets for N≥3 biological replicates. **D.** Cartoon schematic and representative images showing *in vitro* RNA-only droplet assay where *CLN3, BNI1*, and *SPA2* mRNAs assemble into liquid or gel-like droplets. Scale bar 5 μm. **E.** Fluorescence microscopy images showing *BNI1* RNA (green) colocalizes with SPA2 RNA (pink) in droplets. **F.** Fluorescence microscopy images showing *CLN3* RNA (pink) does not colocalize with *SPA2* (green) and *BNI1* (green) droplets. Scale bar 5 μm. **G.** Quantification of co-localization between *BNI1* and *SPA2, SPA2* and *CLN3*, or *BNI1* and *CLN3* RNAs. NS, not significant, ***, p < 0.001 (Wilcoxon rank-sum test). n≥200 droplets for N≥3 biological replicates.

mRNA sequences could influence droplet identity by favoring homotypic or specific heterotypic interactions between RNA molecules. To test for specific RNA-RNA interactions, we used a protein-free system to induce electrostatic-mediated phase transitions of the mRNA (*14*), where all mRNAs were capable of homotypic assembly into liquid or gel-like droplets (Fig. 2D). Strikingly, *CLN3* mRNAs had minimal co-localization with *BNI1* or *SPA2* mRNAs, whereas *BNI1* and *SPA2* were significantly more co-localized (Fig. 2 E–G). Thus, sequence-encoded features of the mRNA can underpin the assembly of distinct, immiscible structures.

We next investigated which features of the mRNA sequence generate specificity. An mRNA with scrambled *CLN3* coding sequence (*cln3 scr*) with intact Whi3-binding sites formed Whi3 droplets (Fig. S1E), but no longer showed specificity (Fig. 3, A and C). As the length, nucleotide composition, and Whi3 binding sites were identical, we hypothesized the secondary structure could promote specificity. *CLN3* mRNA heated to 95°C to disrupt secondary structure also readily incorporated into Whi3-*BNI1* droplets (Fig. 3, A and C). Melted *CLN3* mRNA that was slowly refolded (*CLN3* refold) showed significantly less recruitment than melted, but more than native *CLN3* (Fig. 3A and C). Mixing between melted *CLN3* and melted *BNI1* occurred in the presence of Whi3 and in RNA-only reactions, suggesting mixing is initiated by RNA-RNA interactions (Fig. S2). Thus, specificity information in *CLN3* mRNA can be eliminated by disrupting secondary structure.

**Figure 3.**
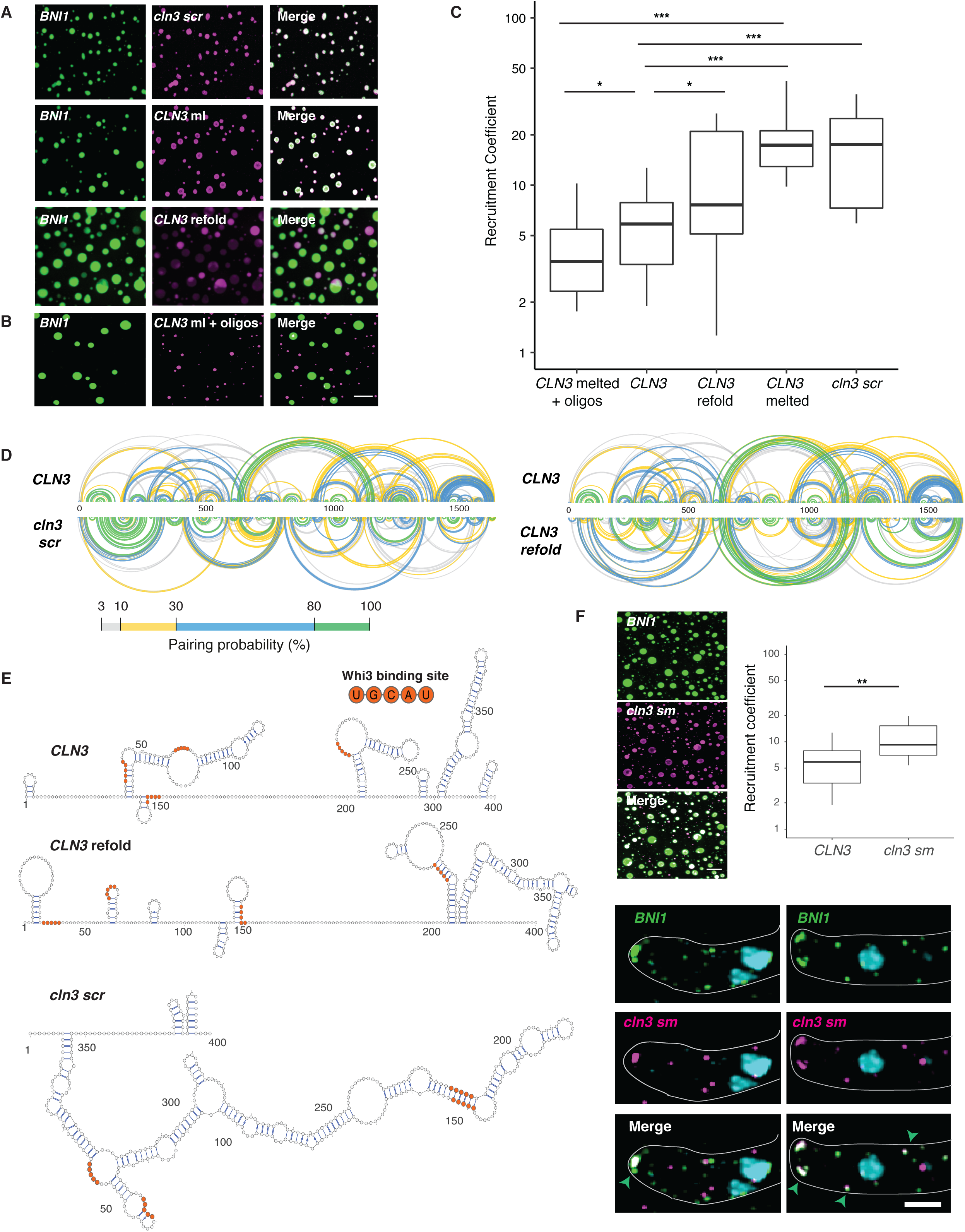
RNA secondary structure determines specificity and identity of Whi3-*CLN3* droplets. **A.** Fluorescence microscopy images showing the recruitment of scrambled (*cln3 scr*), melted (*CLN3* ml), and refolded *CLN3* (*CLN3* refold) mRNA (pink) into preformed Whi3-*BNI1* droplets (green). **B.** Fluorescence microscopy images showing the loss of recruitment of *CLN3* ml when mixed with oligonucleotides targeting complementary sequences of *CLN3* to *BNI1*. Scale bar 10 μm. **C.** Quantification of **A** and **B**. *, p<0.05; **, p<0.01; ***, p < 0.001 (t test). n≥500 droplets for N≥3 biological replicates. **D.** Base pairing probability from SHAPE-MaP of *CLN3, cln3 scr*, and *CLN3* refold show differences in the secondary structure in *CLN3*. Arcs connect base pairs and are colored by probability. **E.** Secondary structure models from SHAPE-MaP for the first 400 nucleotides of *CLN3, CLN3* refold, and *cln3 scr*. Whi3 binding sites are in orange. **F.** *CLN3* structure mutant (*cln3 sm*) mRNA is significantly recruited to Whi3-*BNI1* droplets *in vitro* and *in vivo*. **, p<0.01 (t test). Green arrows denote sites of co-localization between *BNI1* mRNA (green) and *cln3 sm* mRNA (pink) by smFISH. Scale bar 10 μm for *in vitro*, 2 μm for *in vivo*. n ≥500 droplets for N≥3 biological replicates.

To identify what features of *CLN3* mRNA secondary structure promote specificity, we performed SHAPE-MaP, which identifies highly flexible regions in RNA (*15*), to determine secondary structure changes, on native, refolded, and scrambled *CLN3* mRNA (Fig. 3D, S3A and B). The first 400 nucleotides in the *CLN3* sequence exhibited especially low SHAPE reactivity (Fig. S3C, purple shaded regions), suggesting many paired nucleotides and a highly folded structure. Refolded *CLN3* had a significant increase in SHAPE reactivity compared to native *CLN3* (Fig. S3A, p <0.001, Wilcoxon rank sum test), indicating a transition to a more unstructured state (Fig 3D and E). Melting and refolding thus allows the RNA to sample different conformations from those formed during transcription. As expected, *cln3 scr* showed a different SHAPE profile with dramatically altered secondary structure (Fig. 3D and E, S3B).

We hypothesized secondary structure influences mRNA sorting, as stem-loops may selectively display or mask sequences capable of hybridizing with other RNAs. *CLN3* contains five complementary regions to *BNI1* (Fig. S4A), most of which had low SHAPE reactivity and therefore were more structured (Fig. S4B), suggesting these regions are inaccessible for hybridizing with *BNI1*. We hypothesize these regions became available to pair with *BNI1* when *CLN3* is melted, causing the structure-dependent loss of droplet specificity. To test this hypothesis, oligonucleotides (*i.e.*, oligos) complementary to these regions were added to melted *CLN3* and significantly decreased the co-assembly with *BNI1*, restoring the formation of distinct *CLN3* droplets (Fig 3B and C). Additionally, *cln3sm*, a mutant perturbing structure and exposing complementarity, co-localized with *BNI1* transcripts in vitro and at polarity sites in cells (Fig. 3F, S5, >60% tips co-localized). Thus, secondary structure can regulate RNA sorting into distinct droplets through altering the capacity to form intermolecular interactions.

We next asked if exposed complementarity explains co-assembly of *BNI1* and *SPA2* into the same droplets. Indeed, SHAPE-MaP analysis of *BNI1* and *SPA2* showed complementary regions between these co-localizing mRNAs having significantly higher SHAPE reactivity and less structure compared to the *CLN3*/*BNI1* regions (Fig S4 and S6; p < 0.002, t test). Addition of complementary oligos to these regions disrupted co-localization in the presence of Whi3 and in RNA-only reactions (Fig. S7 A and B). We predicted that *CLN3* may self-assemble and indeed *cln3 codon*, a *CLN3* mutant whose codons have been randomized but Whi3 binding sites remain intact, does not co-localize with endogenous *CLN3* mRNA in cells, further supporting RNA-RNA interactions in co-assembly of related RNAs (Fig S7C). These data suggest RNA-RNA interactions based on intermolecular hybridization direct RNAs into the same or different droplets.

Does Whi3 protein influence the identity of droplets? The majority of Whi3 binding sites are exposed on stem loops in *CLN3, BNI1*, and *SPA2* (Fig. 3E, S8 and S9). Notably, refolding or scrambling the *CLN3* sequence rearranges the presentation of Whi3 binding sites (Fig. 3E). Therefore, RNA secondary structure may influence Whi3 binding and contribute to droplet composition and immiscibility in addition to RNA complexing. SHAPE-MaP of *CLN3* mRNA in the presence of Whi3 support that Whi3 binding sites are occupied by protein (Fig. 4A, S10A) and revealed that protein binding causes structural rearrangements (Fig. 4B). We therefore hypothesize Whi3 binding may have important contributions to structural rearrangements of target RNAs relevant to droplet identity.

**Figure 4.**
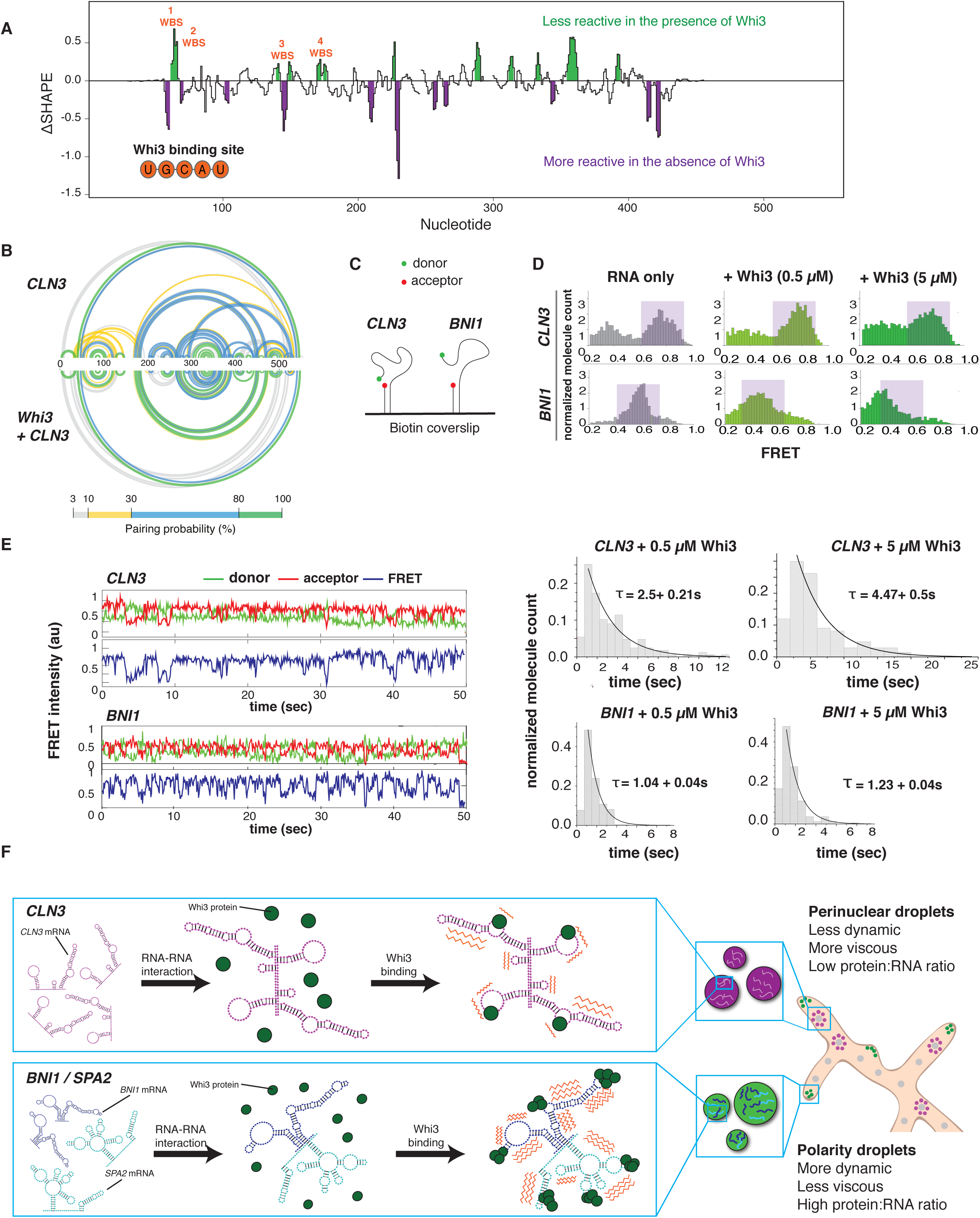
Whi3 binding alters RNA behavior. **A.** Differences in SHAPE reactivities (δSHAPE) were calculated by subtracting *CLN3* SHAPE reactivities from *CLN3* + Whi3 reactivities. Positive δSHAPE values indicate protection from modification in the presence of Whi3 and negative δSHAPE reports enhanced reactivity in the absence of Whi3 protein. **B.** Base pairing probability compared between *CLN3* and *CLN3* with Whi3 shows rearrangements in *CLN3* structure in the presence of Whi3. Arcs connect base pairing sites and are colored by probability. **C.** Schematic of smFRET experiment. **D.** FRET histograms before (gray) and after (green) 0.5 or 5 μM Whi3 addition. Purple shaded regions denote high and mid FRET states for *CLN3* and *BNI1*, respectively. **E.** Averaged cy3 (green), cy5 (red) intensities, and representative FRET traces (blue) obtained from smFRET experiments of *CLN3* and *BNI1* in the presence of 5 μM Whi3. Dwell time analysis reveals slower FRET fluctuations for *CLN3* than *BNI1* in the presence of Whi3. **F.** Proposed model in which RNA-RNA interactions derived from mRNA structure promotes the selective uptake of distinct RNAs and protein constituents into droplets leading to distinct dynamics (orange zigzags) of different droplet complexes.

To examine the consequence of Whi3 binding to RNA, we used smFRET (Fig. 4C) to measure the conformational dynamics of *CLN3* and *BNI1* mRNAs with and without Whi3 (*16*). In the absence of protein, *CLN3* RNA showed high FRET values indicative of a compacted state, while *BNI1* RNA showed lower FRET values, indicating a less compact state (Fig. 4D, purple shaded regions). Upon addition of Whi3, *CLN3* FRET values decreased, indicating more extended RNA conformations were induced, dependent on the ability of Whi3 to bind mRNA (Fig S10 B and C). In contrast, bound to Whi3, *BNI1* RNA showed a more substantial broadening of FRET values (Fig. 4D), indicating Whi3-*BNI1* complexes are more dynamic. Dwell-time analysis revealed Whi3-induced dynamics are three times faster for *BNI1* than *CLN3* (Fig. 4E). Different mRNAs thus react differentially in their intramolecular fluctuations to the presence of Whi3, providing an additional mode of RNA droplet regulation.

These FRET studies suggest Whi3 binding alters the conformational dynamics of target RNAs. We speculate these differential dynamics help maintain droplet identities established by RNA-RNA interactions. Once RNA-RNA interactions are formed, Whi3 binding may reduce the ability of the RNA to resample many alternate RNA structures to maintain the identity. Additionally, the slower fluctuations of *CLN3* bound to Whi3 may be one source of exclusion from the more rapidly fluctuating *BNI1-*Whi3 complexes in those droplets. Such dynamics may drive the droplet material properties reported previously (*3*) and serve as barriers to homogenization.

We show mRNA structure defines the ability of an RNA to engage in homo-or heteromeric interactions and thus drives specificity in the composition of liquid droplet compartments (Fig. 4F). This mechanism is likely relevant for the sorting of specific RNAs to other RNA-granules such as stress and P granules, and P-bodies (*17*, *18*). Future work will address the timing and location of how mRNA secondary structure influences selective uptake of cellular constituents into droplets. Protein binding to different RNAs can lead to varied dynamics of complexes, further distinguishing the physical properties of different droplets and promoting immiscibility of coexisting droplets. Given the large number of distinct, RNA-based condensates in the cell, these mechanisms are likely broadly relevant to explain how droplets achieve and maintain individuality.

## Acknowledgments

We thank the Gladfelter, Weeks, and Laederach labs for critical discussions, Drs Griffin, Moseley, Lew, Peifer, and Higgs for critically reading the manuscript, the HHMI HCIA at the Marine Biological Laboratory for intellectual community, and Timothy Straub for useful data analysis discussions.

## Funding

This work was supported by NIH GM R01-GM081506, the HHMI Faculty Scholars program, R35 GM122532, ACS 130845-RSG-17-114-01-RMC, NIH 1DP2 GM105453, and NIH R01 GM115631.

## Author contributions

EML and ASG designed and performed experiments, analyzed data, prepared figures, and drafted the manuscript; PB, AGN, and CW designed and performed experiments, analyzed data, and edited manuscript; GAM and CMT performed experiments and analyzed data; TMG, JAS, and JMC provided technical support and edited manuscript; KMW and SM designed experiments and edited manuscript.

## Competing interests: Disclosure

K.M.W. is an advisor to and holds equity in Ribometrix, to which mutational profiling (MaP) technologies have been licensed. All other authors declare that they have no competing interests.

## Data and materials availability

All data is available upon request from EML or ASG.

